# High resolution long-read telomere sequencing reveals dynamic mechanisms in aging and cancer

**DOI:** 10.1101/2023.11.28.569082

**Authors:** Tobias T. Schmidt, Carly Tyer, Preeyesh Rughani, Candy Haggblom, Jeffrey R. Jones, Xiaoguang Dai, Kelly A. Frazer, Fred H. Gage, Sissel Juul, Scott Hickey, Jan Karlseder

## Abstract

Telomeres are the protective nucleoprotein structures at the end of linear eukaryotic chromosomes. Telomeres’ repetitive nature and length have traditionally challenged the precise assessment of the composition and length of individual human telomeres. Here, we present Telo-seq to resolve bulk, chromosome arm-specific and allele-specific human telomere lengths using Oxford Nanopore Technologies’ native long-read sequencing. Telo-seq resolves telomere shortening in five population doubling increments and reveals intrasample, chromosome arm-specific, allele-specific telomere length heterogeneity. Telo-seq can reliably discriminate between telomerase- and ALT-positive cancer cell lines. Thus, Telo-seq is a novel tool to study telomere biology during development, aging, and cancer at unprecedented resolution.

## Main text

Mammalian telomeres, the nucleoprotein structures at the end of eukaryotic linear chromosomes, consist of 5’-TTAGGG-3’ repeats and terminate in a single-stranded G-rich overhang ^1, 2^. The overhang can fold back and form a telomeric loop (T-loop)^3^. The hexameric protein complex shelterin binds telomeric repeats and stabilizes the T-loop ^4^. Together, the T-loop ^5, 6^ and shelterin ^4, 7^ protect the telomeric chromosome ends from being recognized as DNA double-strand breaks ^8^. Due to the “end-replication problem” and subsequent processing, telomeres in human somatic cells shorten with every round of DNA replication ^1, 9, 10^. Short, deprotected telomeres are recognized by the DNA damage response and either trigger a permanent cell cycle arrest named replicative senescence ^11^ or, in cells deficient for the p53 and Rb checkpoint pathways, replicative crisis ^12–14^, a state with extensive innate immunity-driven, autophagy-dependent cell death ^15, 16^. Both replicative senescence and crisis restrict the maximum number of cell divisions of human somatic cells and act as powerful, telomere-dependent proliferation barriers against human carcinogenesis ^17^. Thus, to overcome proliferation barriers and acquire replicative immortality, cancer cells must activate a telomere maintenance mechanism (TMM) ^18^. Whereas most cancers reactivate the reverse transcriptase telomerase (TERT) ^19, 20^, 10-15% of all cancers maintain their telomeres by the recombination-based “Alternative Lengthening of Telomeres” (ALT) mechanism ^21^. The absence of TMM in human somatic cells makes TMM-specific vulnerabilities attractive targets for personalized cancer therapy ^22^; however, it is critical to efficiently distinguish between ALT^+^ and TERT^+^ cancers, which is currently challenging in the clinical setting.

To measure telomere length, various methods have been developed, including Terminal Restriction Fragment (TRF) analysis ^9^, STELA ^23^, TeSLA ^24^, quantitative PCR ^25^, Q-FISH ^26^, flow FISH ^27, 28^, DNA combing ^29^ and telomere length estimates based on next-generation sequencing data ^30^. These methods use either telomere enrichment, staining of telomeres with specific probes, or a combination of both. However, these traditional methods fail to resolve chromosome arm and allele-specific composition of individual telomeres due to their repetitive nature and length. With the advent of DNA long-read sequencing, it is now possible to sequence entire telomeres and harvest subtelomeric information to annotate individual telomeric reads to specific chromosome arms. In the budding yeast *Saccharomyces cerevisiae*, chromosome arm-specific telomere length and telomere shortening was resolved using Oxford Nanopore Technologies long-read sequencing ^31^. Further, nanopore long-read sequencing was recently applied to measure telomere shortening in an RTEL1 mutant mouse model and to compare mouse telomere length to the human one^32^. For human telomeres, a recent report combined a telomeric pulldown with restriction enzyme digest and PacBio HiFi long-read sequencing to measure telomere length and telomere variant repeats in cultured human cells and patient cells ^33^. However, due to the protocol’s stringent restriction enzyme digest, the annotation of telomeric reads to chromosome arms was only possible for nine chromosome arms, and the very long telomeres present in ALT^+^ cells ^21^ are incompatible with the processivity of the PacBio HiFi DNA polymerase ^33^. Furthermore, data from the Telomere-to-Telomere (T2T) ^34, 35^ and Genome in the Bottle Consortiums ^36^ demonstrated that human whole-genome long-read sequencing can be utilized to analyze human telomere length and composition. However, as the telomeric content of human diploid cells is approximately only 0.015% of the total genome, telomere length measurements based on whole-genome long-read sequencing are inefficient and telomere enrichment is necessary. Here, we developed Telo-seq to efficiently sequence entire human telomeres using nanopore sequencing and applied it to explore bulk, chromosome arm and allele-specific human telomere length and composition in aging and cancer.

### Telo-seq

To enrich for telomeres, telorette-based telomere adapters ^23, 24^ were first annealed to the G-overhang and ligated to the C-strand (Fig. 1a). Next, genomic DNA was digested with the blunt-end restriction enzyme EcoRV. To reduce concatemer ligation, a dA-tail was added to the blunt ends prior to splint adapter annealing and sequencing adapter ligation. After nanopore sequencing, bases were called using a customized Bonito telomere model (Extended Data Fig. 1a). Next, the reads were filtered for quality, the telomeric motif was identified and reads filtered for expected structure (Extended Data Fig. 1a,b). To annotate reads to individual chromosome arms, reads were mapped to a collection of well-annotated subtelomeric sequences ^34, 37^ (Extended Data Fig. 1c).

**Fig. 1.**
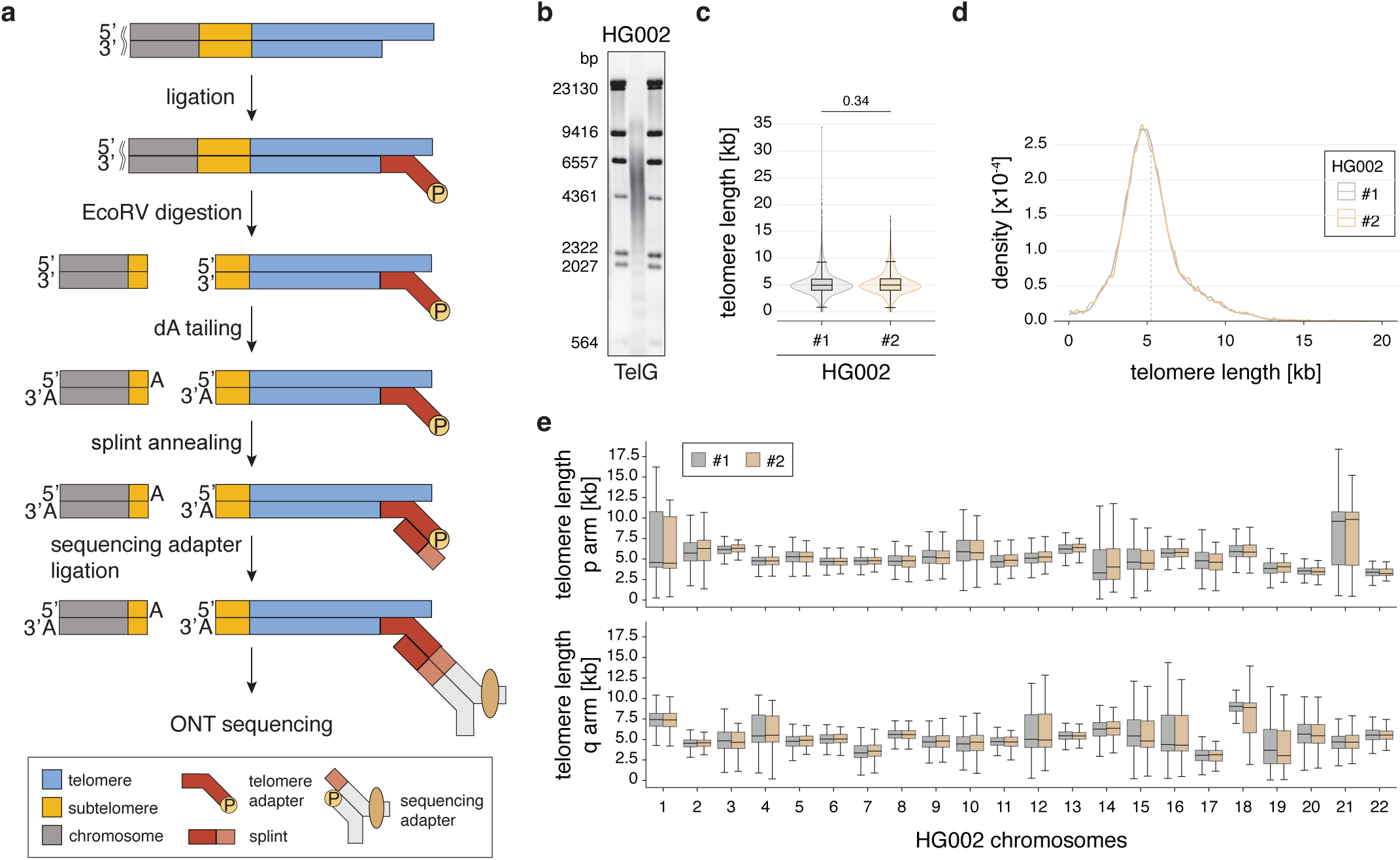
Telo-seq measures bulk and chromosome arm specific telomere length. **a**, Schematic overview of Telo-seq protocol. **b**, Terminal Restriction Fragment (TRF) analysis of HG002. **c**, Violin plot of HG002 Telo-seq telomere length measurements. Violin represents telomere length distribution in kilobases (kb). Boxplot shows the median telomere length with interquartile range (IQR) and whiskers represent 1.5-fold IQR. Statistical analysis t-test with Bonferroni correction was performed and adjusted p-value is shown. **d**, HG002 Telo-seq telomere length distribution with the mean telomere length shown as the dotted line. **e**, Boxplot of the chromosome arm-specific telomere lengths of both HG002 replicates. The middle line represents the median telomere length, box the IQR and whiskers 1.5-fold IQR. For (c-e), results of two independent HG002 replicates are shown.

To evaluate Telo-seq, we used the B-lymphocyte cell line HG002, which was sequenced previously as part of the Human Pangenome Reference and Genome in a Bottle Consortiums and a high-quality telomere-to-telomere assembly is publicly available ^37, 38^. First, we compared telomere enrichments of Telo-seq to long-read sequencing of high-molecular weight DNA and AluI/MboI double-digested genomic DNA (the restriction enzymes used in traditional TRF analysis ^16, 39^). Telo-seq resulted in most telomeric reads of all tested protocols and increased telomeric reads 75-fold relative to whole-genome sequencing (Extended Data Fig. 1d). Second, we compared Telo-seq bulk telomere length measurement of two independent HG002 replicates with TRF. Based on TRF analysis, HG002 cells have a bulk telomere length of 4 to 6 kilobases (kb) (Fig. 1b). In line with the TRF results, Telo-seq revealed a mean telomere length of 5,247 and 5,270 base pairs (bp) for each independent replicate, with a standard deviation of 2,085 and 2,135 bp, respectively (Fig. 1c,d, Supplementary Table 1). Furthermore, both replicates showed very similar subtelomere and telomere length distributions (Fig. 1c,d, Extended Data Fig. 1e, Supplementary Table 1). Finally, to assess chromosome arm-specific telomere length, individual reads were anchored to chromosome arms using the subtelomeric sequences (Extended Data Fig. 1c). The normalized coverage of mapped telomeric reads per chromosome arm revealed consistent and uniform coverage across replicates (Extended Data Fig. 1f). The intrasample telomere lengths between different chromosome arms were heterogenous, ranging in the two replicates from median telomere lengths of 3,088 and 3,139 bp at chromosome 17q to 9,603 and 9,835 bp at chromosome 21p (Fig. 1e). However, despite the intrasample heterogeneity, the chromosome arm-specific telomere lengths were highly reproducible in the two replicates (Extended Data Fig. 1g). We therefore conclude that Telo-seq can reproducibly measure bulk and chromosome arm-specific telomere lengths of human cells.

### Telo-seq resolves telomere shortening

In the absence of an active TMM, telomeres progressively shorten due to the end-replication problem and subsequent processing ^1, 9, 10^. To address whether Telo-seq can resolve telomere shortening dynamics, IMR90 human lung fibroblasts expressing the human papilloma virus E6 and E7 oncogenes (IMR90 ^E6E7^) were grown *in vitro* to replicative crisis and sampled at different population doublings (PD) for telomere length analysis ^16, 17^. In line with the absence of a TMM, TRF analysis revealed that IMR90 ^E6E7^ telomeres progressively shorten with increasing PDs (Fig. 2a). Similarly, telomere shortening was also detected by Telo-seq (Fig. 2b-d, Extended Data Fig. 2a, Supplementary Table 2). The IMR90 ^E6E7^ mean bulk telomere length shortened from 4,276 bp at PD66.2 to 2,746 bp at PD106.1. The fraction of short telomeres below 1 kb in length was progressively increasing from 5.0% at PD66.2 to 18.3% at PD106.1, while the fraction of telomeres above 10 kb decreased from 3.2% to 1.0% (Fig. 2d, Supplementary Table 2). By plotting the mean telomere length against the PDs and performing linear regression analysis (Fig. 2e), we estimated that IMR90 ^E6E7^ telomeres shorten on average around 39 bp per PD under the growth conditions used in our laboratory.

**Fig. 2.**
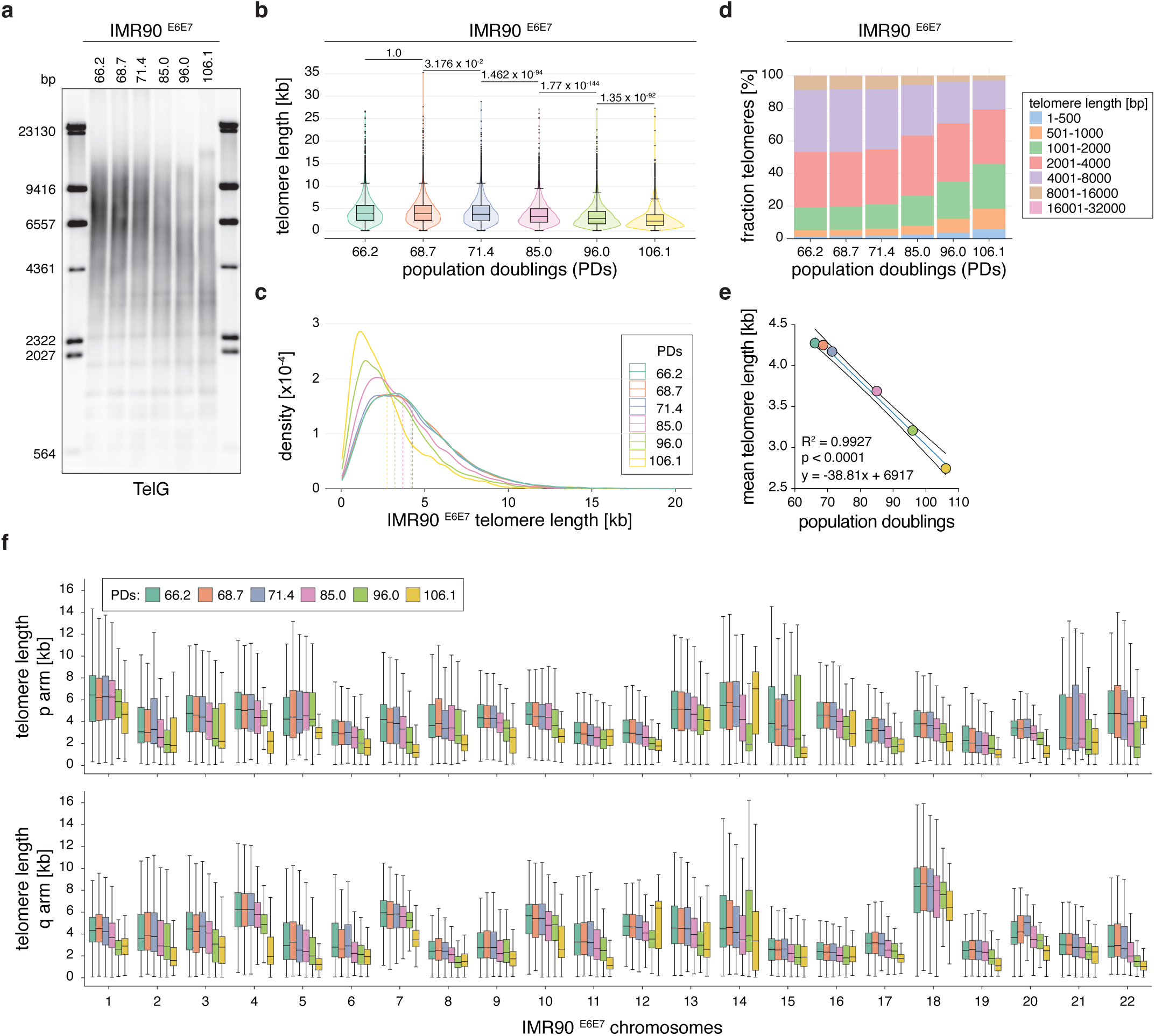
Telo-seq resolves telomere shortening. **a,** Terminal Restriction Fragment (TRF) analysis of human IMR90 ^E6E7^ fibroblasts at different population doublings (PDs). **b**, Violin plot of IMR90 ^E6E7^ Telo-seq telomere length measurements in kilobases (kb) at indicated PDs. Violin represents telomere length distribution. Boxplot shows the median telomere length with interquartile range (IQR) and whiskers represent 1.5-fold IQR. Statistical analysis t-test with Bonferroni correction was performed and adjusted p-values are shown. **c,** IMR90 ^E6E7^ Telo-seq telomere length distribution at different PDs with the mean telomere length shown as the dotted line. **d**, Bar graph showing the percentage of binned telomere length in IMR90 ^E6E7^ fibroblasts at different PDs. **e**, Linear regression analysis of IMR90 ^E6E7^ mean telomere length against PDs. Blue line represents best fit and black curves 95% confidence intervals. **f,** Boxplot of chromosome arm specific IMR90 ^E6E7^ telomere length at different PDs. The middle line represents the median telomere length, box the IQR and whiskers 1.5-fold IQR.

Next, we analyzed IMR90 ^E6E7^ chromosome arm-specific telomere length dynamics (Fig. 2f, Extended Data Fig. 2b,c). Similar to the bulk telomere length analysis, IMR90 ^E6E7^ telomeres of individual chromosome arms progressively shortened with increasing PDs, but at variable shortening rates (Fig. 2f, Extended Data Fig. 2c). As indicated by the HG002 chromosome arm-specific telomere length analysis (Fig. 1e), telomere lengths between different chromosome arms in IMR90 ^E6E7^ were highly heterogenous (Fig. 2f). For example, at PD 66.2 the median telomere length of chromosome 18q was 8,350 bp, whereas the median telomere at chromosome 19q was 2,275 bp long. Taken together, Telo-seq can measure bulk and chromosome arm-specific telomere length dynamics and resolve telomere shortening rates of samples only five PDs apart.

### Telo-seq on donor-derived fibroblasts revealed shorter telomeres with human age

To investigate telomere length in aged human individuals, we next performed Telo-seq on patient-derived fibroblasts obtained from donors between 20 to 94 years of age. Telomeres were generally longer in the younger individuals than in the older individuals, except for an 82-year-old individual who harbored telomeres of comparable length to the young individuals (Fig. 3a,b, Extended Data Fig. 3a, Supplementary Table 3). In line with that, the fraction of short telomeres below 1 kb and long telomeres above 10 kb in length were lowest and highest in the young individuals, respectively (Supplementary Table 3). Plotting of the mean telomere length against the donor age and linear regression analysis revealed a general trend of bulk telomere shortening as function of donor age (Fig. 3c). Telomere length assessment of individual chromosome arms confirmed the intrasample heterogeneity of chromosome arm-specific telomere length observed in HG002 and IMR90 ^E6E7^ (Extended Data Fig. 3b,c). Thus, we asked whether any chromosome arms consistently possessed shorter or longer telomeres than the average telomere length in the sample. To address this, we combined the eight fibroblast samples with the IMR90 ^E6E7^ PD66.2 sample and ranked chromosome arms according to their mean telomere length. Based on these 9 individuals and despite the heterogeneity, some chromosome arms consistently had shorter or longer telomeres, relative to the mean telomere length (Extended Data Fig. 3d), raising the possibility that conserved chromosome arm-specific features influence telomere length.

**Fig. 3.**
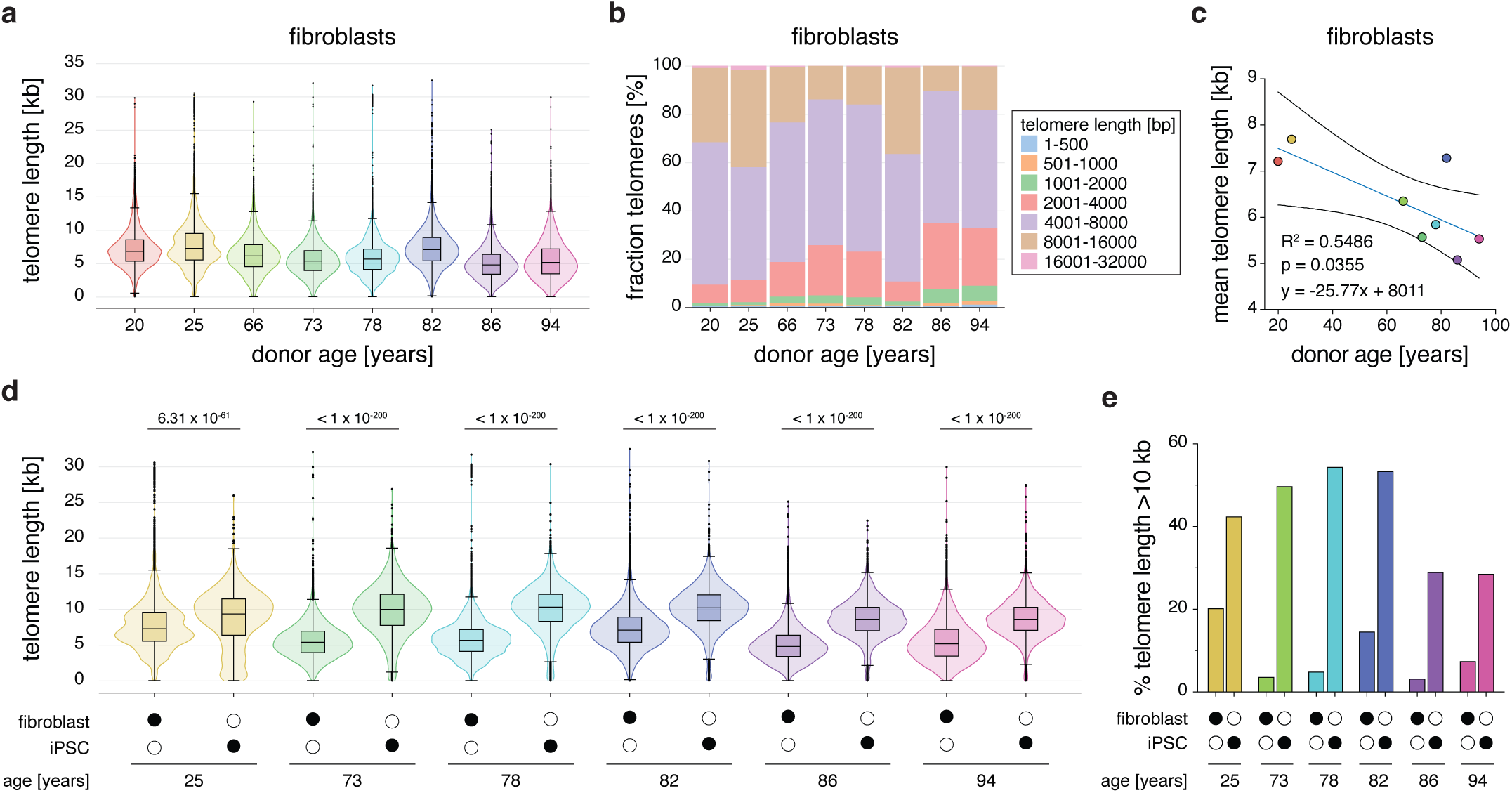
Telomeres shorten with age. **a,** Violin plots of donor-derived fibroblast Telo-seq telomere length measurements in kilobases (kb). Violin represents telomere length distribution. Boxplot shows the median telomere length with interquartile range (IQR) and whiskers represent 1.5-fold IQR. **b,** Bar graph showing the percentage of binned telomere length in doner-derived fibroblasts. **c,** Linear regression analysis of fibroblast mean telomere length against donor age. Blue line represents best fit and black curves 95% confidence intervals. **d,** Violin plots of donor-derived fibroblast and matched induced pluripotent stem cells (iPSC) Telo-seq telomere length measurements. Violin represents telomere length distribution. Boxplot shows the median telomere length with IQR and whiskers represent 1.5-fold IQR. Statistical analysis t-test with Bonferroni correction was performed and adjusted p-values are shown. **e,** Bar graph showing the percentage of telomeres longer than 10 kb of donor-derived fibroblasts and their matched iPSC.

### Telomere length differs between alleles

Our analysis suggested that bulk telomere length heterogeneity is partly a consequence of the chromosome arm-specific telomere length heterogeneity. We speculated that, similarly to the bulk telomere length, the intrachromosome arm-specific telomere length heterogeneity could partially originate from the two alleles of each chromosome arm. To test this, we first used HG002 to resolve allele-specific telomere length by mapping the reads to the phased HG002 reference genome (Extended Data Fig. 4a). Indeed, some of the intrachromosome arm-specific heterogeneity could be explained by differences in the allele-specific telomere length. For example, the HG002 chromosome arm 1p maternal allele had a median telomere length of 4,169 bp, whereas the median paternal allele was 11,158 bp long (Fig. 1e, Extended Data Fig. 4a). As the mapping approach requires a phased reference genome for a given sample, we tested whether allele-specific telomere length can also be quantified by de-novo haplotype phasing (Extended Data Fig. 4b). Indeed, both approaches retrieved comparable allele-specific telomere length information for HG002. However, the strict implementation of phasing introduced certain limitations, resulting in the absence of alignments to some chromosomes and alleles. This can be attributed to two key factors. Firstly, conventional phasing algorithms are designed solely for primary alignments and do not consider secondary alignments. Secondly, the use of a custom pangenome reference results in higher quality alignments compared to a haploid reference. We applied the de-novo haplotype phasing approach to IMR90 ^E6E7^ PD66.2 in order to resolve IMR90^E6E7^ allele-specific telomere length (Extended Data Fig. 4c). Similar to HG002, the IMR90 ^E6E7^ allele-specific telomere length was frequently more homogenous than at the chromosome arm level (Extended Data Fig. 4c). Thus, in addition to bulk and chromosome arm-specific telomere length assessment, Telo-seq allows the analysis of higher resolution allele-specific telomere length.

### Telomere length is reset in donor-matched iPSCs

Upon induction of pluripotency, aging hallmarks ^40^ are reversed, including telomere shortening, which is counteracted by telomerase ^41–44^. To investigate the effect of induced pluripotency on telomere length, we performed Telo-seq on six matched induced pluripotent stem cells (iPSCs) of fibroblast donors between 25 and 94 years of age (Fig. 3d, Extended Data Fig. 5a,b, Supplementary Table 4). Telomeres in iPSCs were on average 3,202 bp longer than in their matched primary fibroblasts. Furthermore, the fraction of telomeres longer than 10 kb was increased 2- to 12-fold in iPSCs relative to their matched fibroblast controls (Fig. 3e). The dispersion in the telomere length distribution was reduced in iPSCs suggesting that telomerase activity results in more homogenous telomere length (Supplementary Table 3, 4). Independently of the telomere length in matched fibroblasts and the donor’s age, iPSCs mean telomeres were set to 8 to 10 kb (Fig. 3d, Supplementary Table 3, 4), indicating that in this range human iPSCs’ telomeres are in an equilibrium, regulated by telomerase-dependent elongation and telomere trimming ^45^.

Based on linear regression analysis of chromosome arm-specific mean telomere lengths with at least a 20x coverage in matched fibroblasts and iPSCs (Extended Data Fig. 5c-f), we concluded that upon induced pluripotency shorter telomeres are preferentially elongated by telomerase; however, the overall order of chromosome arm-specific telomere length remained conserved between fibroblasts and iPSCs.

### Telo-seq distinguishes between TERT^+^ and ALT^+^ cancer cells

Cancer cells maintain their telomeres by either reactivation of telomerase or ALT ^19–21^. To address the impact of both TMMs on telomeres, we performed Telo-seq on a set of five TERT^+^ and five ALT^+^ cancer cell lines. TRF analysis revealed that bulk telomere length in TERT^+^ cell lines was more tightly distributed, whereas telomeres of ALT^+^ cancer cell lines were more heterogenous in length (Fig. 4a)^46^. Telo-seq analysis recapitulated this striking difference in the telomere length distribution between TERT^+^ and ALT^+^ cancer cell lines (Fig. 4b,c, Supplementary Table 5) and was highly reproducible between independent cancer cell line replicates (Supplementary Table 6). In ALT^+^ cells, we could measure telomeres from 47 bp to 134.7 kb in length (Supplementary Table 5) suggesting that Telo-seq is an efficient approach to resolve the very long telomeres present in ALT^+^ cells. Next, we speculated whether Telo-seq can distinguish between ALT^+^ and TERT^+^ cancer cells by plotting the coefficient of variation (CV), a measure of the dispersion in a distribution, against mean telomere length (Fig 4d). Indeed, all ALT^+^ cancer cells had a CV above 0.8, whereas the TERT^+^ cancer cells showed a CV smaller than 0.55, indicating that Telo-seq is an effective method to postulate TMM of cancer cells.

**Fig. 4.**
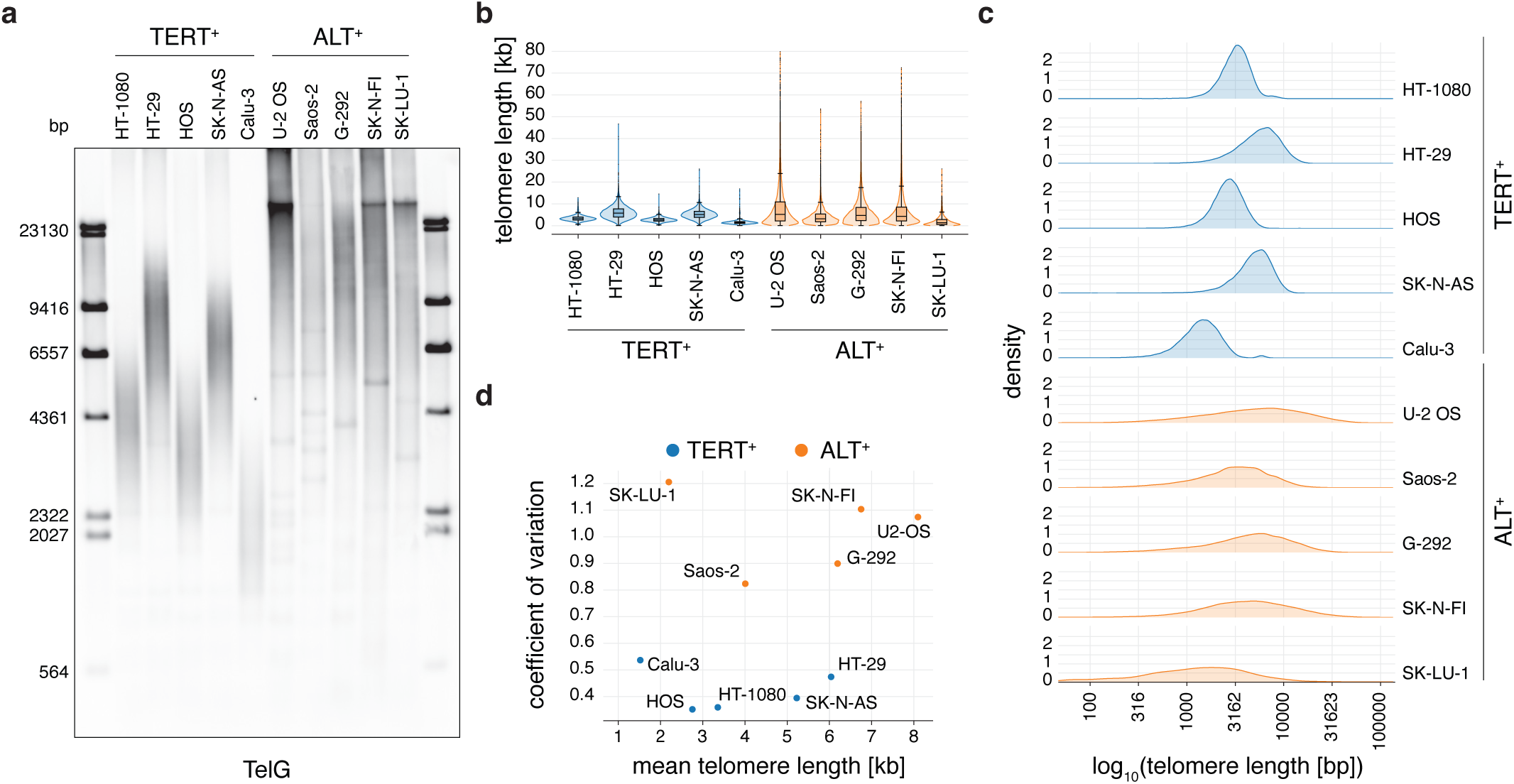
Telo-seq resolves TERT^+^ and ALT^+^ cancer cell telomere length distributions. **a**, Terminal Restriction Fragment (TRF) analysis of the indicated cancer cell lines. **b,** Violin plot of cancer cell line Telo-seq telomere length measurements of telomeres shorter than 80 kilobases (kb). Violin represents telomere length distribution. Boxplot shows the median telomere length with interquartile range (IQR) and whiskers represent 1.5-fold IQR. **c**, Density plot of telomere length distributions measured by Telo-seq in indicated cancer cell lines. Telomere length in base pairs (bp) is log10 transformed. **d,** Scatter plot of coefficient of variations to mean telomere length in kb of indicated cancer cell lines.

The subtelomeres of most human chromosome arms contain a CpG island adjacent to the telomere. As Telo-seq uses native DNA, we retrieved the subtelomeric DNA methylation information from the long nanopore reads. In line with previous studies ^47^, the subtelomeric CpG islands were frequently hypomethylated in ALT^+^ cancer cell lines (Extended Data Fig. 6, 7). Not all the chromosome arms were equally hypomethylated and we found differences within and between ALT^+^ cancer cell lines. Thus, Telo-seq allows not only telomere length measurements, but also chromosome arm-specific subtelomeric methylation analysis.

## Discussion

Here, we developed Telo-seq, an efficient and reproducible method to determine the length and sequence of entire human telomeres and part of the adjacent subtelomere using Oxford Nanopore Technologies (Fig. 1a). We demonstrate that Telo-seq resolves the short telomeres present in crisis cells ^48^ to the very long telomeres existing in ALT^+^ cancer cell lines ^21^. We find that intrasample telomere length is heterogenous and short telomeres are identified even in cells many population doublings away from proliferation barriers (Fig. 2, Supplementary Table 2). Harvesting subtelomeric information enabled us to assess chromosome arm and allele-specific telomere length (Fig. 1e, 2f, Extended Data Fig. 3c, 4, 5d). In line with previous work quantifying chromosome arm-specific telomeric FISH staining on metaphases ^49^ and nanopore sequencing of budding yeast telomeres ^31^, our work shows that within one sample chromosome arms, and even alleles, vary in their telomere length distributions. Hence, some of the detected bulk and chromosome arm-specific length heterogeneity can be explained by the chromosome arm and allele-specific telomere length variability, respectively. Furthermore, our analysis suggests that some of these chromosome arm-specific telomere length differences are conserved between different individuals (Extended Data Fig. 3d). This may indicate that like in budding yeast ^31^, there are conserved chromosome arm-specific factors that influence telomere length homeostasis in humans that are yet to be identified. Telo-seq analysis of a large human cohort will define these conserved chromosome arms better and, in combination with genome editing and high content imaging, will serve to identify mechanisms as to why these chromosome arms consistently differ from the average telomere length.

Telomere shortening is a hallmark of aging ^40^ and an elegant protective mechanism to prevent infinite proliferation ^17^. In line with previous telomere shortening rates for human fibroblasts ^9, 50^, we find that the mean telomere length in IMR90 ^E6E7^ fibroblasts shorten by approximately 39 bp per PD (Fig. 2e). Interestingly, individual chromosome arms vary in their shortening rate (Extended Data Fig. 2c), likely pointing at chromosome arm-specific factors that influence telomeric processing ^1^. Similarly, telomere length and attrition may vary in different tissues and cell types of the same individual depending on tissue-specific differential gene expression, the cellular turnover rate, the exposure to endogenous and exogenous sources of DNA damage, as well as inflammation ^51^. Telo-seq on tissues or sorted cell types will therefore not only reveal precise telomere length measurements, but also provide information about telomeric sequence composition and subtelomeric methylation status during human aging and disease.

Consistent with previous reports ^42–44^, we show that upon induction of pluripotency the mean telomere length is reset independently of the parental cell’s initial telomere length (Fig 3d). Our results indicate that telomerase preferentially elongates shorter telomeres (Extended Data Extended Data Fig. 5e), which not only leads to increased bulk telomere length (Fig. 3d), but also a more homogenous telomere length distribution (Supplementary Table 3, 4). Moreover, our analysis of cancer cells reveals that Telo-seq can efficiently distinguish between TERT^+^ and ALT^+^ cancer cells (Fig. 4b-d), establishing Telo-seq as reliable method of TMM prediction, an essential prerequisite to target TMM-specific vulnerabilities in a personalized cancer therapy ^22^. In summary, our study highlights the potential of human telomere long-read sequencing and sets the stage to investigate human telomere dynamics in unprecedented detail during development, aging and disease.

## Methods

### Cell culture

Calu-3 (HTB-55), G-292, clone A141B1 (CRL-1423), HOS (CRL-1543), HT-1080 (CCL-121), HT-29 (HTB-38), Saos-2 (HTB85), SK-LU-1 (HTB-57), SK-N-AS (CRL-2137), SK-N-FI (CRL-2142) and U-2 OS (HTB96) cancer cell lines and IMR90 (CCL-186) fibroblasts were purchased from ATCC. The lymphoblastoid cell line HG002 (GM24385) was purchased from Coriell Institute for Medical Research. Donor-derived fibroblasts were collected at University of California, San Diego (UCSD) and are part of the Salk AHA-Allen aging cohort. Alzheimer’s Disease Research Center participants at UCSD have given broad consent to a range of experiments, including skin fibroblast and induced pluripotent stem cell derivation, cell engineering, and genetic sequencing and manipulation prior to providing a skin biopsy.

All cells were grown at 37 °C under 7.5% CO_2_ and 3% O_2,_ except HG002 and induced pluripotent stem cells (iPSCs) were grown at 37 °C under 5% CO_2_ and ambient O_2_. IMR90 fibroblasts were grown in GlutaMax-DMEM (Gibco, 10569-010) supplemented with 0.1 mM non-essential amino acids (Corning, 25-025-Cl), 15% fetal bovine serum (FBS) (VWR, 97068-085 Lot 323B20), 100 IU/mL Penicillin and 100 μg/ml Streptomycin (Corning, 30-002-CI). Donor-derived fibroblasts were grown in GlutaMax-DMEM supplemented with 0.1 mM non-essential amino acids, 20% FBS, 20 ng/mL FGF-2 (Joint Protein Central), 100 IU/mL Penicillin and 100 μg/ml Streptomycin. Calu-3, HOS, HT-1080, Saos-2, SK-LU-1, SK-N-AS, SK-N-FI, U-2 OS were grown in GlutaMax-DMEM supplemented with 0.1 mM non-essential amino acids, 10% FBS, 100 IU/mL Penicillin and 100 μg/ml Streptomycin. HT-29 and G-292 were grown in McCoy-s 5a (modified) media (Gibco, 16600-108) supplemented with 10% FBS and 100 IU/mL Penicillin and 100 μg/ml Streptomycin. HG002 cells were grown in Roswell Park Memorial Institute 1640 medium (ATCC, 30-2001) and 10% FBS (ATCC, 30-2020) and 1% antibiotic antimycotic (Gibco, 15240062). iPSCs were grown in StemMACS™ iPS-Brew XF, human (Miltenyi, 130-104-368) with daily medium replacement.

The number of population doublings (PD) of IMR90 fibroblasts were calculated using the following equation: PD = log(cells harvested / cells seeded)/log2. Cells have been tested to be free of mycoplasma.

### Induced pluripotent stem cells generation

Induced pluripotent stem cells (iPSCs) were generated from donor-derived dermal fibroblasts at the Salk stem cell core according to standards of the International Society for Stem Cell Research standards ^52^. In brief, dermal fibroblasts were expanded and verified mycoplasm negative via MycoAlert™ PLUS Mycoplasma Detection Kit (Lonza, 75860-362) infected with Sendai virus containing Yamanaka factors from the CytoTune™-iPS 2.0 Sendai Reprogramming Kit (ThermoFisher, A165167) according to manufacturer recommendations. iPSCs were grown in medium sized colonies on Matrigel (BD biosciences, 354230) at a final concentration of 1 mg per 6-well plate and propagated in StemMACS™ iPS-Brew XF, human (Miltenyi, 130-104-368) with daily medium replacement.

### Terminal Restriction Fragment analysis

Terminal Restriction Fragment (TRF) was performed as previously described ^15, 53^. In brief, high-molecular weight genomic DNA (gDNA) was isolated by phenol-chloroform extraction and digested with 50 U AluI (NEB, R0137L) and 50 U MboI (NEB, R0147M) at 37 °C, overnight. Digested gDNA was quantified using Qubit dsDNA BR assay (Invitrogen, Q32850) and either 4 or 5 μg digested gDNA was separated on a 0.7% agarose gel overnight in 1x TAE buffer at 40 V. Next, the gel was incubated in depurination buffer (0.25 M HCl) for 10 min followed by two 15 min incubations in denaturing buffer (1.5 M NaCl, 0.5 M NaOH) and two 15 min incubations in neutralization buffer (1 M Tris, pH 7.4, 1.5 M NaCl). The gDNA was transferred to a positively charged nylon membrane (Amersham, RPN203B), overnight. After UV-crosslinking, the membrane was incubated in prehybridization buffer (5x SSC, 0.1% N-lauroylsarcosine sodium salt and 0.04% sodium dodecyl sulfate (SDS)) for 2 h at 65 °C followed by hybridization overnight at 65 °C (1.3 nM digoxigenin-labeled TelG probe prepared according to ^54^ in prehybridization buffer). The next day, the membrane was washed with wash buffer 1 (2x SSC + 0.1% SDS) for 15 min, thrice, followed by one 15 min wash in wash buffer 2 (2x SSC). The membrane was blocked with freshly prepared blocking solution (100 mM maleic acid, 150 mM NaCl, pH 7.5, 1% (wt/vol) blocking reagent (Roche, 11096176001)) for 30 min. Next, the membrane was incubated with an anti-digoxigenin-AP antibody (Roche, 11093274910, 1:2000 dilution in blocking solution) for 30 min. Membrane was washed twice with wash buffer 3 (1x maleic acid buffer, 0.3% Tween 20) for 15 min and equilibrated with AP buffer (100 mM Tris, 100 mM NaCl, pH 9.5) for 2 min. Blot was developed using CDP-star ready to use solution (Roche, 12041677001) and imaged in G-box (Syngene).

### Genomic DNA Extraction and Purification for Oxford Nanopore Technologies sequencing

Genomic DNA was extracted from 5-10×10^6^ cells using the Gentra Puregene Cell Kit (Qiagen, cat no. 158046) following the manufacturer’s instructions. Extracted DNA was further purified by isopropanol precipitation. DNA was quantified using the Qubit fluorometer (Thermo Fisher).

### Telo-seq

Six 5’-phosphorylated single-stranded oligo adapters containing permutations of an 18 bp sequence complementary to the human telomeric repeat were combined in equal parts to a final concentration of 1 µM. Oligo sequences are shown in Supplementary Table 7. A separate splint oligonucleotide S1 with sequence complementarity to the Oxford Nanopore AMII sequencing adapter was diluted to a final concentration of 10 µM. The pre-mixed oligos were ligated with T4 DNA ligase (NEB, M0202) using the following reaction conditions, as previously described in ^24^: 20 µL pre-mixed oligos, 10,000 U T4 DNA ligase, 10 mM ATP (NEB, P0756), 10x rCutSmart Buffer, and 15 µg extracted DNA in a final volume of 200 µL. The ligation mixture was incubated at 35 °C for 16 hours and then heat inactivated at 65 °C for 10 min. Adapter-ligated DNA was digested with 4 µL of EcoRV-HF (NEB, R3195), incubated at 37 °C for 30 minutes, and heat inactivated at 65 °C for 20 min. Digested DNA was treated with Klenow Fragment (3’→5’ exo-) (NEB, M0212) in a final volume of 250 µL of 1x NEBNext dA-tailing reaction buffer (NEB), split evenly into 5 aliquots, and incubated at 37 °C for 30 min. The resulting DNA was purified using 1x v/v AMPure XP (Beckman Coulter, A63881) and annealed at 50 °C for 1 hour in the presence of 50 mM NaCl and 100 nM splint oligo. Final DNA was purified using 0.5x v/v AMPure XP and sequencing libraries were prepared with the AMII sequencing adapter following the Native Barcoding Kit instructions (Oxford Nanopore Technologies, SQK-NBD111). Libraries were sequenced on a GridION sequencer (Oxford Nanopore Technologies) with R9.4.1 flow cells.

### Analysis

#### Telomere Identification and Length Determination

Reads were basecalled using a telomere Bonito basecalling model trained for calling telomeric repeats. Identification of telomere-containing reads was achieved through the implementation of the Noise Cancelling Repeat Finder (NCRF) algorithm (version 1.01.00 20190426)^55^. Subsequent reads were further filtered based on three criteria:

1. Due to the nature of Oxford Nanopore sequencing reads from the 5’ to 3’ direction, reads that did not commence with the 5’-(CCCTAA)_n_ motif or conclude with the 5’-(TTAGGG)_n_ motif were excluded from the analysis.
2. Identification of the telomere motif within 200 base pairs of the read termini.
3. The inclusion of 500 bp unique subtelomeric sequence adjacent to the telomere motif.

Telomere length determination was carried out by utilizing the outcomes obtained from NCRF. In cases where a read contained multiple segments due to telomeric variations spanning over 50 base pairs, these segments were concatenated into a unified sequence, provided that the breakpoints between segments did not exceed 250 base pairs. This concatenated sequence was then considered a single, uninterrupted length measurement for the telomere.

#### Read Mapping and Alignment

All filtered reads were mapped against a custom reference containing “Telomere-to-Telomere” (T2T) consortium reference genomes HG002 (v0.7)^34^ and CHM13 (v2.0)^38^. For HG002, we retained 25 kilobases on each end of the chromosomes for both maternal and paternal copies; for CHM13, we retained the entire genome. Using Minimap2 (Version 2.22), with specific alignment parameters optimized for nanopore reads (minimap -x map-ont -N 2 -Y -y -L -a). Alignments were ranked following a hierarchical order of primary, secondary, and supplementary alignments. In scenarios where the primary alignment lacked intersection with both subtelomeric or telomeric regions, secondary alignments were examined iteratively until the intersection criteria were fulfilled. In cases where secondary alignments met the above criteria, they were promoted to primary status for compatibility with downstream analyses, and the original primary alignments were subsequently reclassified as secondary.

#### Haplotyping

Fully phased reads were obtained by mapping telomere reads from HG002 to the phased diploid reference HG002(v0.7) ^34^ using Minimap2. Maternal and paternal reads were obtained based on their primary alignment to the contig, filtering for reads with a mapping quality greater than 5 (Extended Data Fig. 4a).

For samples where a phased reference genome is not available, a de-novo haplotyping approach to quantify allele-specific telomere lengths was developed (Extended Data Fig. 4b,c). Telomere reads were aligned using Minimap2 to the paternal haploid copy of the reference genome HG002 (v0.7) ^34^ and variants were called using Clair3with the r941_hac_g360_g422_1235 model. The resulting reads were haplotagged with WhatsHap haplotag.

We cross-referenced haplotagged reads with the prior alignments obtained using the custom pangenome reference and only considered reads that matched precisely on read id, chromosome, arm, and strand (see Extended Data Fig. 4b,c). Phased reads with primary alignments to the haploid reference are shown; secondary alignments were not considered.

#### Methylation

Telomeric reads were basecalled with Dorado 0.3.4 using the dna_r9.4.1_e8_sup@v3.3_5mC_5hmC model. The reads were aligned to a custom pangenome reference using Minimap2. Methylation calls were obtained using Modkit (v0.11.1) pileup with the “traditional” preset. A minimum coverage threshold of 10 reads was applied to the final output.

#### Statistical analysis

The statistical analysis was conducted using GraphPad Prism (version 8.4.3), R (version 4.3.1) and Python (version 3.8.10). In Python, the telomere length distributions were compared using the t-test of independence with Bonferroni correction. For ranking chromosomes, the z-score was calculated with the Python library Scipy.

## Acknowledgments

This manuscript was submitted back-to-back with one from the Artandi lab (S. E. Sanchez et al.). We thank S. Artandi and S.E. Sanchez for exchanging information and coordinating the manuscripts.

We thank the entire Karlseder lab for discussions. This work was supported by the Stem Cell Core Facility of the Salk Institute with funding from the Helmsley Charitable Trust, the Shiley-Marcos Alzheimer’s Disease Research Center (ADRC; AG062429) at the University of California, San Diego (UCSD), and the National Institute of Aging of the National Institutes of Health (P30AG068635). This research was supported by an AHA-Allen Initiative in Brain Health and Cognitive Impairment award made jointly through the American Heart Association and The Paul G. Allen Frontiers Group: 19PABH134610000.

---

**Supplementary Table 1.**
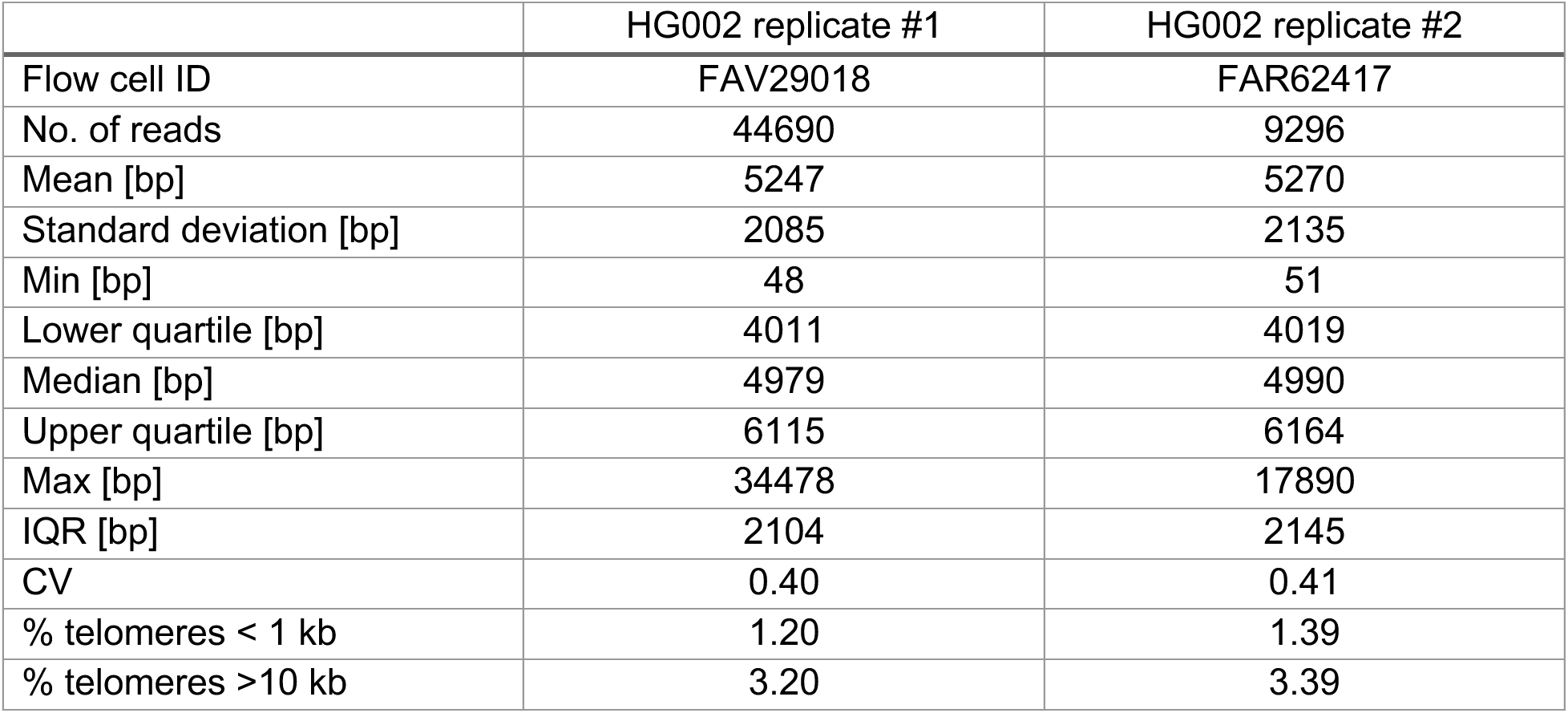
Telo-seq summary HG002.

**Supplementary Table 2.** Telo-seq summary IMR90 ^E6E7^ progression.

|  | IMR90<sup>E6E7</sup> |  |  |  |  |  |
| --- | --- | --- | --- | --- | --- | --- |
| Population doublings | 66.2 | 68.7 | 71.4 | 85.0 | 96.0 | 106.1 |
| Flow cell ID | FAU74032 | FAU64354 | FAV31378 | FAW19571 | FAW29975 | FAV35454 |
| No. of reads | 32301 | 23243 | 18699 | 27469 | 33409 | 13135 |
| Mean [bp] | 4276 | 4252 | 4176 | 3691 | 3211 | 2746 |
| Standard deviation [bp] | 2610 | 2606 | 2597 | 2381 | 2219 | 2161 |
| Min [bp] | 48 | 50 | 49 | 50 | 49 | 51 |
| Lower quartile [bp] | 2359 | 2340 | 2246 | 1932 | 1561 | 1221 |
| Median [bp] | 3808 | 3811 | 3713 | 3223 | 2732 | 2176 |
| Upper quartile [bp] | 5668 | 5650 | 5599 | 4945 | 4325 | 3587 |
| Max [bp] | 26695 | 35280 | 28771 | 27151 | 27148 | 27340 |
| IQR [bp] | 3309 | 3310 | 3353 | 3013 | 2764 | 2366 |
| CV | 0.61 | 0.61 | 0.62 | 0.65 | 0.69 | 0.79 |
| % telomeres <1 kb | 4.98 | 5.33 | 5.97 | 7.78 | 11.99 | 18.32 |
| % telomeres >10 kb | 3.24 | 3.08 | 2.96 | 1.73 | 1.13 | 0.99 |

**Supplementary Table 3.**
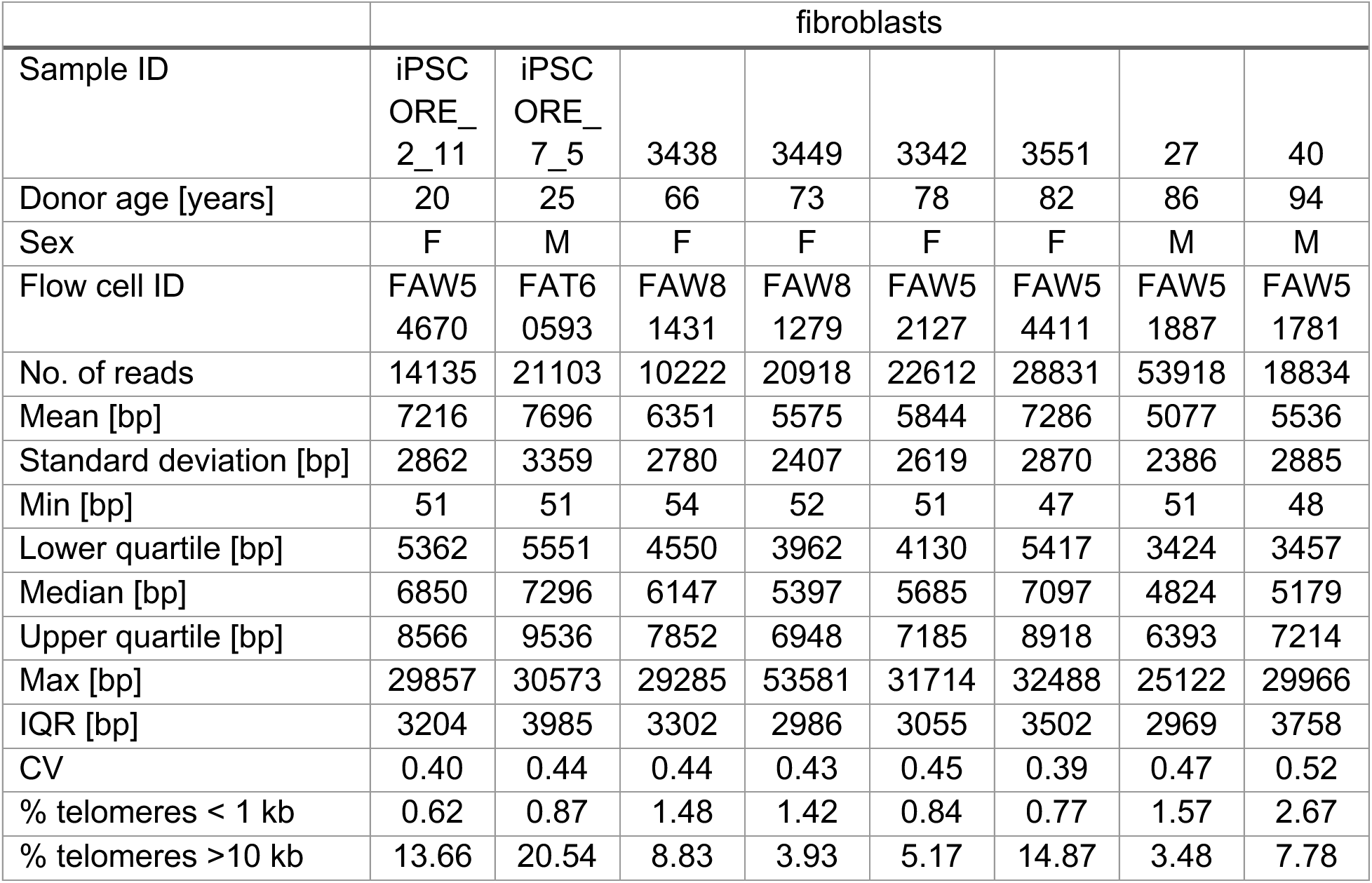
Telo-seq summary fibroblasts of aging cohort.

**Supplementary Table 4.**
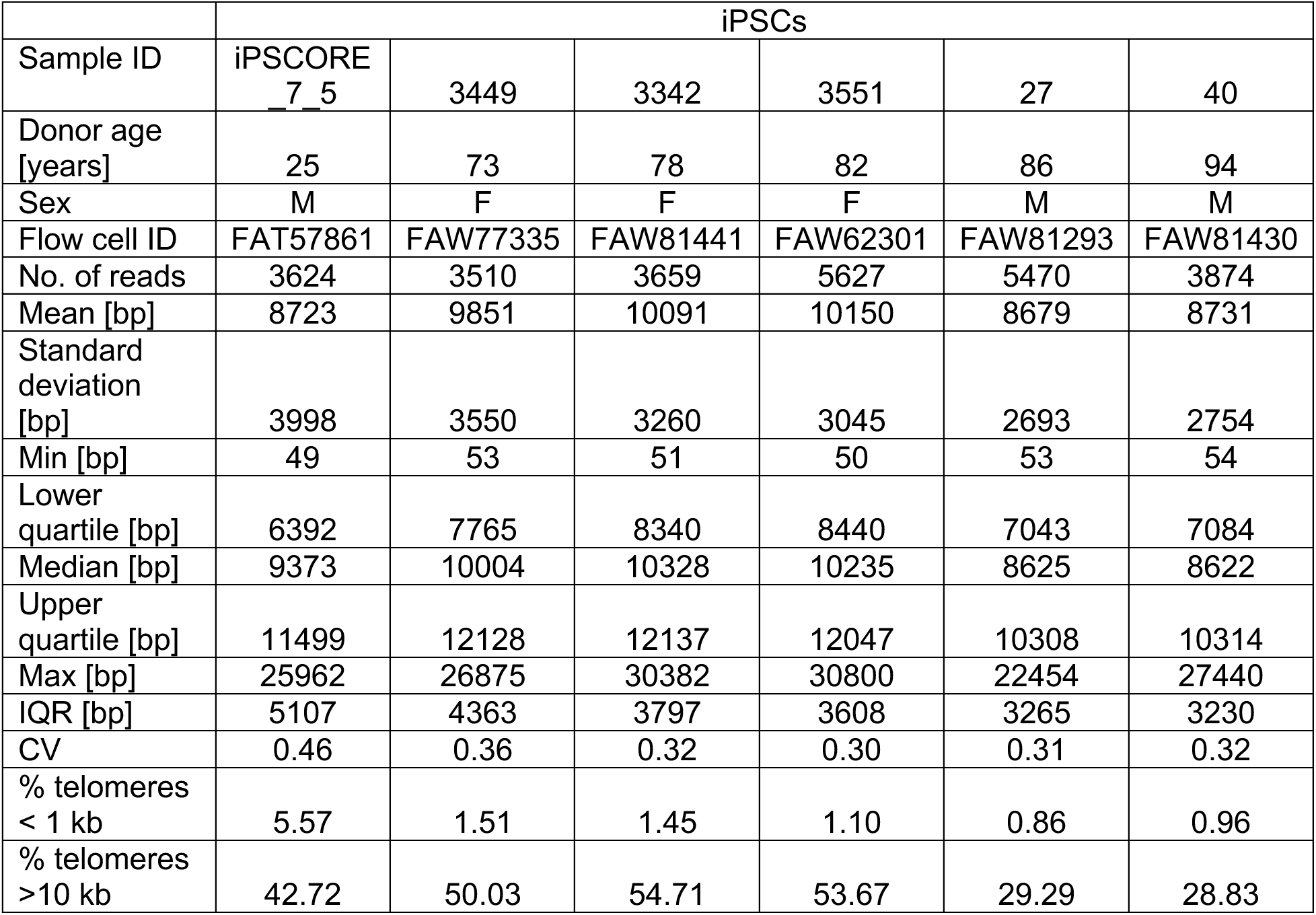
Telo-seq summary iPSCs.

|  | iPSCs |  |  |  |  |  |
| --- | --- | --- | --- | --- | --- | --- |
| Sample ID | iPSCORE<br>7_5 | 3449 | 3342 | 3551 | 27 | 40 |
| Donor age<br>[years] | 25 | 73 | 78 | 82 | 86 | 94 |
| Sex | M | F | F | F | M | M |
| Flow cell ID | FAT57861 | FAW77335 | FAW81441 | FAW62301 | FAW81293 | FAW81430 |
| No. of reads | 3624 | 3510 | 3659 | 5627 | 5470 | 3874 |
| Mean [bp] | 8723 | 9851 | 10091 | 10150 | 8679 | 8731 |
| Standard<br>deviation<br>[bp] | 3998 | 3550 | 3260 | 3045 | 2693 | 2754 |
| Min [bp] | 49 | 53 | 51 | 50 | 53 | 54 |
| Lower<br>quartile [bp] | 6392 | 7765 | 8340 | 8440 | 7043 | 7084 |
| Median [bp] | 9373 | 10004 | 10328 | 10235 | 8625 | 8622 |
| Upper<br>quartile [bp] | 11499 | 12128 | 12137 | 12047 | 10308 | 10314 |
| Max [bp] | 25962 | 26875 | 30382 | 30800 | 22454 | 27440 |
| IQR [bp] | 5107 | 4363 | 3797 | 3608 | 3265 | 3230 |
| CV | 0.46 | 0.36 | 0.32 | 0.30 | 0.31 | 0.32 |
| % telomeres<br>< 1 kb | 5.57 | 1.51 | 1.45 | 1.10 | 0.86 | 0.96 |
| % telomeres<br>>10 kb | 42.72 | 50.03 | 54.71 | 53.67 | 29.29 | 28.83 |

**Supplementary Table 5.** Telo-seq summary cancer cell lines.

|  | HT-1080 | HT-29 | HOS | SK-N-AS | Calu-3 | U2-OS | Saos-2 | G-292 | SK-N-FI | SK-LU-1 |
| --- | --- | --- | --- | --- | --- | --- | --- | --- | --- | --- |
| TMM | TERT<sup>+</sup> | TERT<sup>+</sup> | TERT<sup>+</sup> | TERT<sup>+</sup> | TERT<sup>+</sup> | ALT<sup>+</sup> | ALT<sup>+</sup> | ALT<sup>+</sup> | ALT<sup>+</sup> | ALT<sup>+</sup> |
| No. of flow cells | 1 | 1 | 1 | 2 | 1 | 2 | 2 | 2 | 1 | 4 |
| No. of reads | 20168 | 28512 | 25930 | 50408 | 75064 | 50175 | 65470 | 41308 | 31749 | 1868 |
| Mean [bp] | 3357 | 6040 | 2755 | 5226 | 1522 | 8096 | 4008 | 6193 | 6750 | 2197 |
| Standard deviation [bp] | 1208 | 2864 | 971 | 2063 | 817 | 8694 | 3301 | 5570 | 7448 | 2648 |
| Min [bp] | 50 | 50 | 51 | 47 | 50 | 50 | 49 | 50 | 50 | 49 |
| Lower quartile [bp] | 2582 | 3979 | 2098 | 3790 | 1019 | 2177 | 1732 | 2293 | 2141 | 572 |
| Median [bp] | 3240 | 5733 | 2663 | 5097 | 1406 | 5184 | 3171 | 4688 | 4311 | 1374 |
| Upper quartile [bp] | 3974 | 7719 | 3328 | 6527 | 1866 | 10865 | 5314 | 8381 | 8529 | 2832 |
| Max [bp] | 12853 | 46668 | 14562 | 26026 | 16890 | 134722 | 53547 | 57078 | 108011 | 26053 |
| IQR [bp] | 1392 | 3740 | 1230 | 2737 | 847 | 8688 | 3582 | 6088 | 6388 | 2260 |
| CV | 0.36 | 0.47 | 0.35 | 0.39 | 0.54 | 1.07 | 0.82 | 0.90 | 1.10 | 1.21 |
| % telomeres < 1 kb | 1.01 | 1.23 | 2.11 | 1.05 | 23.84 | 11.00 | 12.51 | 10.39 | 8.65 | 39.78 |
| % telomeres >10 kb | 0.05 | 8.93 | 0.00 | 1.82 | 0.01 | 27.60 | 5.50 | 18.33 | 20.15 | 2.41 |

**Supplementary Table 6.** Telo-seq summary cancer cell lines per flow cell.

| Sample | Flow cell ID | No. of reads | Mean [bp] | Standard deviation [bp] | Min [bp] | Lower quartile [bp] | Median [bp] | Upper quartile [bp] | Max [bp] | IQR [bp] | CV | % telomeres < 1 kb | % telomeres >10 kb |
| --- | --- | --- | --- | --- | --- | --- | --- | --- | --- | --- | --- | --- | --- |
| HT-1080 | FAV27627 | 20168 | 3357 | 1208 | 50 | 2582 | 3240 | 3974 | 12853 | 1392 | 0.36 | 1.01 | 0.05 |
| HT-29 | FAU93874 | 28512 | 6040 | 2864 | 50 | 3979 | 5733 | 7719 | 46668 | 3740 | 0.47 | 1.23 | 8.93 |
| HOS | FAV28723 | 25930 | 2755 | 971 | 51 | 2098 | 2663 | 3328 | 14562 | 1230 | 0.35 | 2.11 | 0.00 |
| SK-N-AS | FAV35348 | 29538 | 5246 | 2061 | 50 | 3805 | 5118 | 6548 | 20460 | 2743 | 0.39 | 0.94 | 1.88 |
| SK-N-AS | FAW30055 | 20870 | 5198 | 2064 | 47 | 3766 | 5074 | 6493 | 26026 | 2727 | 0.40 | 1.19 | 1.73 |
| Calu-3 | FAW31323 | 75064 | 1522 | 817 | 50 | 1019 | 1406 | 1866 | 16890 | 847 | 0.54 | 23.84 | 0.01 |
| U2-OS | FAV89509 | 14493 | 7090 | 7282 | 51 | 2009 | 4722 | 9606 | 134722 | 7597 | 1.03 | 11.93 | 23.78 |
| U2-OS | FAW19540 | 35682 | 8505 | 9175 | 50 | 2253 | 5395 | 11466 | 104101 | 9213 | 1.08 | 10.62 | 29.15 |
| Saos-2 | FAV92737 | 8481 | 3996 | 3307 | 51 | 1723 | 3161 | 5319 | 46266 | 3596 | 0.83 | 12.95 | 5.49 |
| Saos-2 | FAW30039 | 56989 | 4010 | 3301 | 49 | 1734 | 3173 | 5313 | 53547 | 3579 | 0.82 | 12.44 | 5.51 |
| G-292 | FAT60863 | 14621 | 6202 | 5545 | 50 | 2320 | 4720 | 8351 | 56378 | 6031 | 0.89 | 10.23 | 18.28 |
| G-292 | FAW29984 | 26687 | 6189 | 5583 | 50 | 2278 | 4669 | 8393 | 57078 | 6115 | 0.90 | 10.48 | 18.35 |
| SK-N-F1 | FAV34018 | 31749 | 6750 | 7448 | 50 | 2141 | 4311 | 8529 | 108011 | 6388 | 1.10 | 8.65 | 20.15 |
| SK-LU-1 | FAU08605 | 337 | 2072 | 2555 | 50 | 556 | 1333 | 2619 | 23465 | 2063 | 1.23 | 41.54 | 1.78 |
| SK-LU-1 | FAV21836 | 374 | 2035 | 2442 | 52 | 490 | 1218 | 2796 | 20631 | 2307 | 1.20 | 43.32 | 1.87 |
| SK-LU-1 | FAV35378 | 551 | 2268 | 2564 | 51 | 595 | 1435 | 3055 | 20515 | 2460 | 1.13 | 38.48 | 2.54 |
| SK-LU-1 | FAW30048 | 606 | 2302 | 2884 | 49 | 619 | 1434 | 2800 | 26053 | 2181 | 1.25 | 37.79 | 2.97 |

**Supplementary Table 7.**
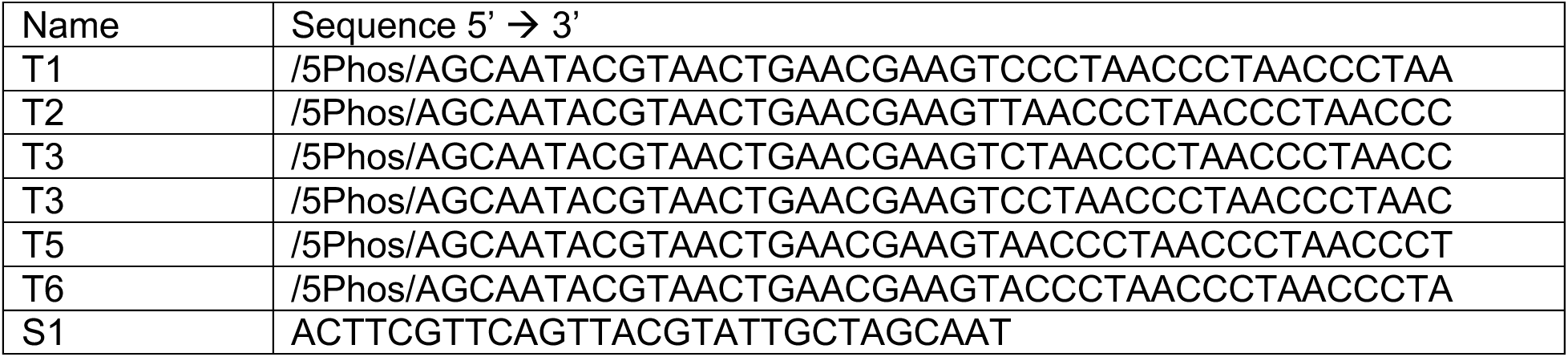
Telo-seq oligos.

## Extended Data Figure legends

**Extended Data Fig. 1.**
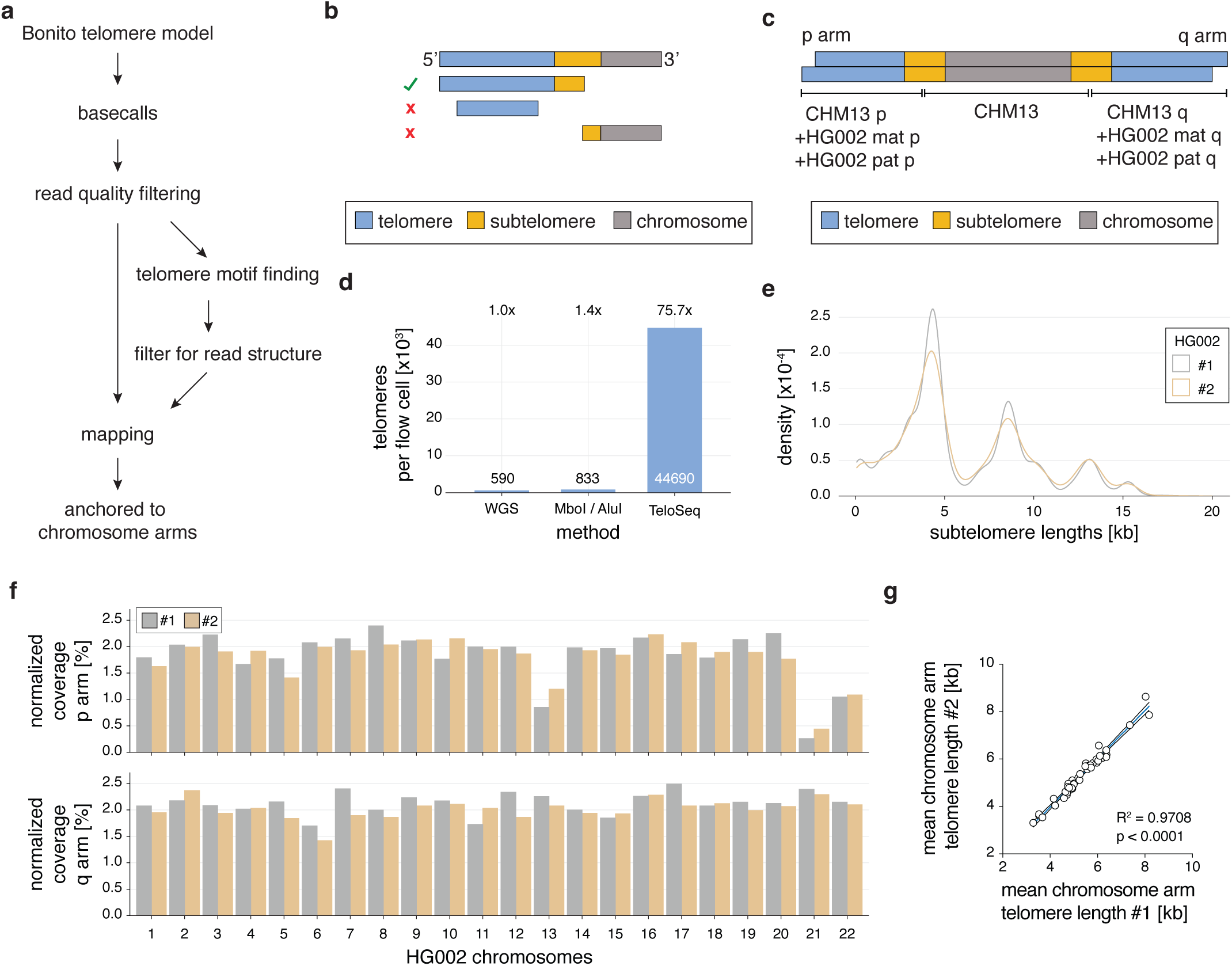
Telo-seq analysis exemplified on HG002. **a,** Schematic overview of bioinformatic analysis of Telo-seq reads. **b,** Schematic representation of the read structure filter. Only reads with a terminal telomeric sequence and an adjacent unique, non-telomeric sequence pass the read structure filter. **c,** Schematic representation of combined assembly used to anchor telomeric reads to specific chromosome arms. CHM13 and HG002 were used for each chromosome arm. **d,** Bar graph comparing the number of telomeric reads per flow cell. Fold increase over whole genome sequencing is given above bar. **e,** Subtelomeric length density for the two independent replicates of HG002. Subtelomeric length is given in kilobase (kb). **f,** Bar graph of the normalized coverage of mapped telomeric reads per chromosome arm for both HG002 replicates in percent. The expected coverage for 46 chromosome arms is ∼ 2.1 % per chromosome arm. **g,** Scatter plot of mean chromosome arm specific telomere length of HG002 replicate 2 against replicate 1. Linear regression analysis was performed. Blue line represents best fit and black curves 95% confidence intervals.

**Extended Data Fig. 2.**
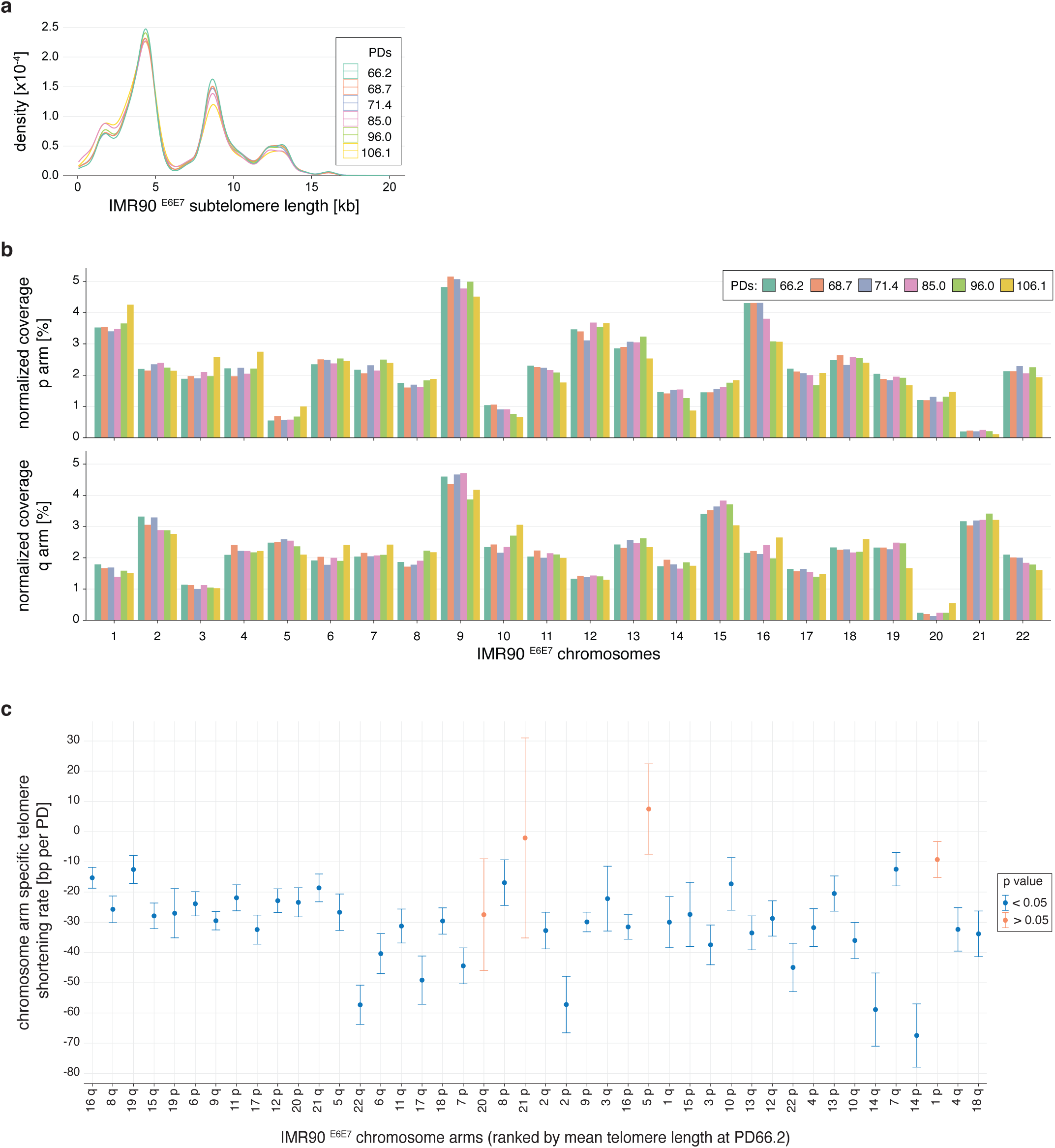
Telo-seq analysis of telomere shortening in IMR90 ^E6E7^ fibroblasts. **a,** Subtelomeric length density of IMR90 ^E6E7^ fibroblasts at different population doublings (PD). Subtelomeric length is given in kilobase (kb). **b,** Bar graph of the normalized coverage of telomeric reads per chromosome arm for IMR90 ^E6E7^ at different PDs in percent. **c,** Chromosome arm-specific telomere shortening rate in base pairs (bp) per PD based on linear regression analysis of chromosome arm-specific telomere length of IMR90 ^E6E7^ from PD66.2 to PD96.0. Rate is shown with standard error. Chromosomes are ranked based on mean telomere length in IMR90 ^E6E7^ at PD66.2.

**Extended Data Fig. 3.**
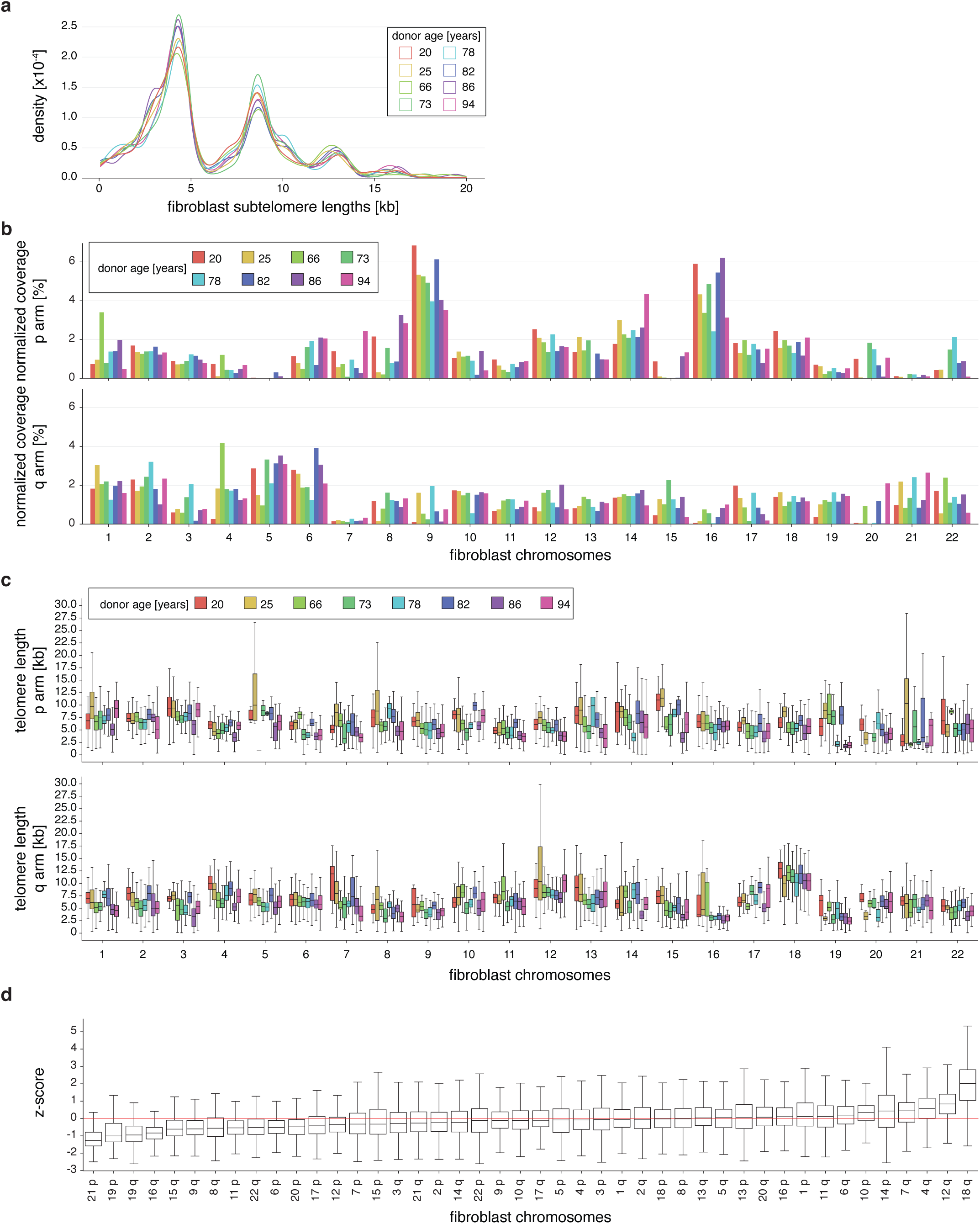
Telo-seq on aging cohort fibroblasts. **a,** Subtelomeric length density of donor-derived fibroblasts of the indicated age. Subtelomeric length is given in kilobase (kb). **b,** Bar graph of the normalized coverage of telomeric reads per chromosome arm of donor-derived fibroblasts of the indicated age in percent. **c,** Boxplot of chromosome arm-specific telomere length of donor-derived fibroblasts with the indicated donor age. The middle line represents median telomere length, box the interquartile range (IQR) and whiskers the 1.5-fold IQR. **d,** z-score analysis of fibroblast chromosome arm-specific telomere length of donor-derived fibroblasts and IMR90 ^E6E7^ PD66.2. The middle line represents median z-score, box the IQR and whiskers the 1.5-fold IQR. Chromosome arms are ranked according to their median z-score. The z-score of the median bulk telomere length of all nine fibroblast samples is shown as red line.

**Fig. S4.**
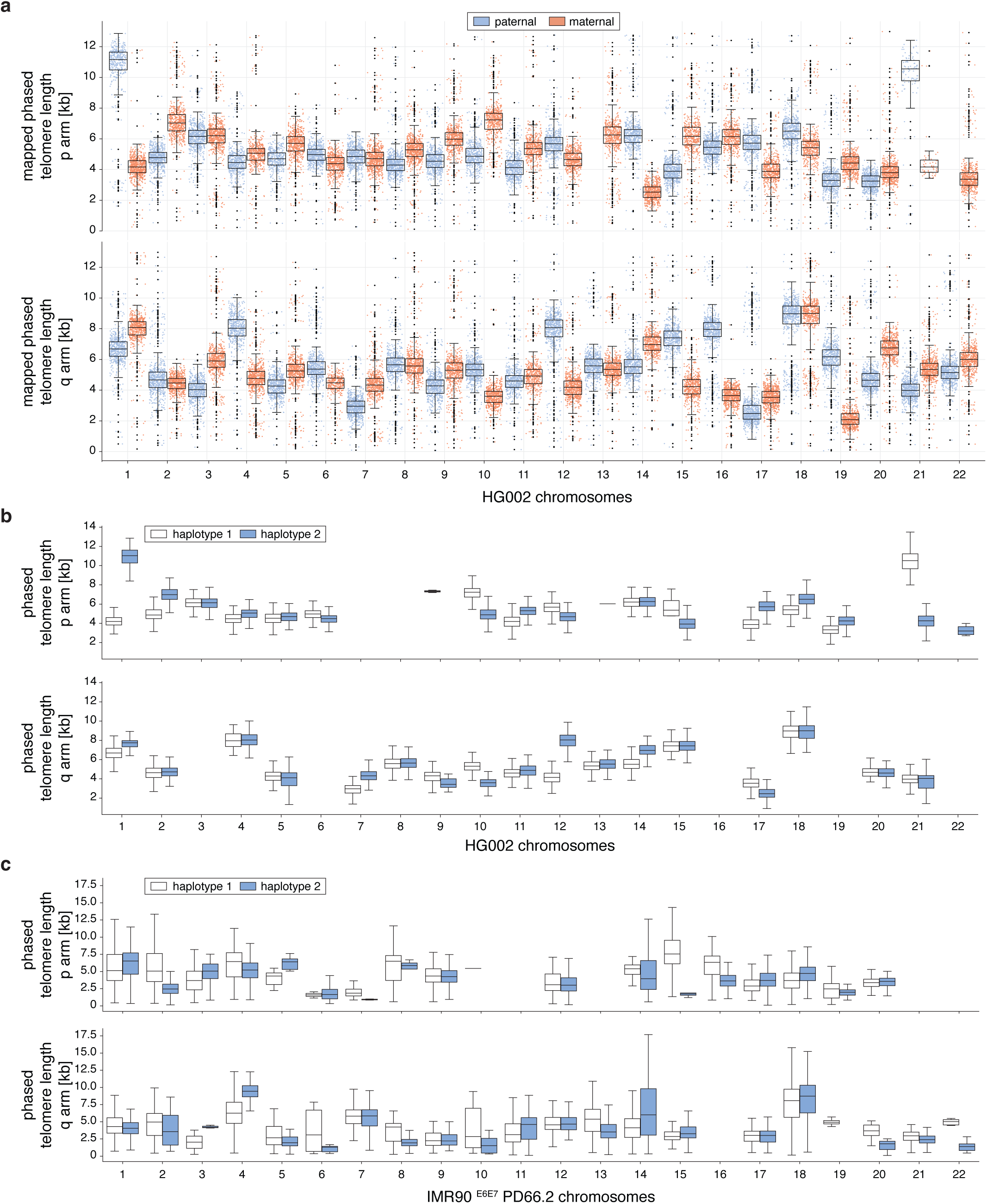
Allele specific telomere length of HG002 and IMR90 ^E6E7^. **a,** Boxplot of allele-specific telomere length in kilobase (kb) of HG002 based on mapping against the HG002 reference genome. The middle line represents median, box the interquartile range (IQR) and whiskers the 1.5-fold IQR with outliers shown as black points. Individual telomere reads are shown as blue and red points. **b,** Boxplot of allele-specific telomere length in kb of HG002 based on de-novo haplotype phasing. The middle line represents median, box the interquartile range (IQR) and whiskers the 1.5-fold IQR. **c,** Boxplot of allele-specific telomere length in kb of IMR90 ^E6E7^ at population doubling (PD) 66.2 based on de-novo haplotype phasing. The middle line represents median, box the interquartile range (IQR) and whiskers the 1.5-fold IQR. For b and c only the alleles that passed the filter were included in the figure, see methods for details.

**Fig. S5.**
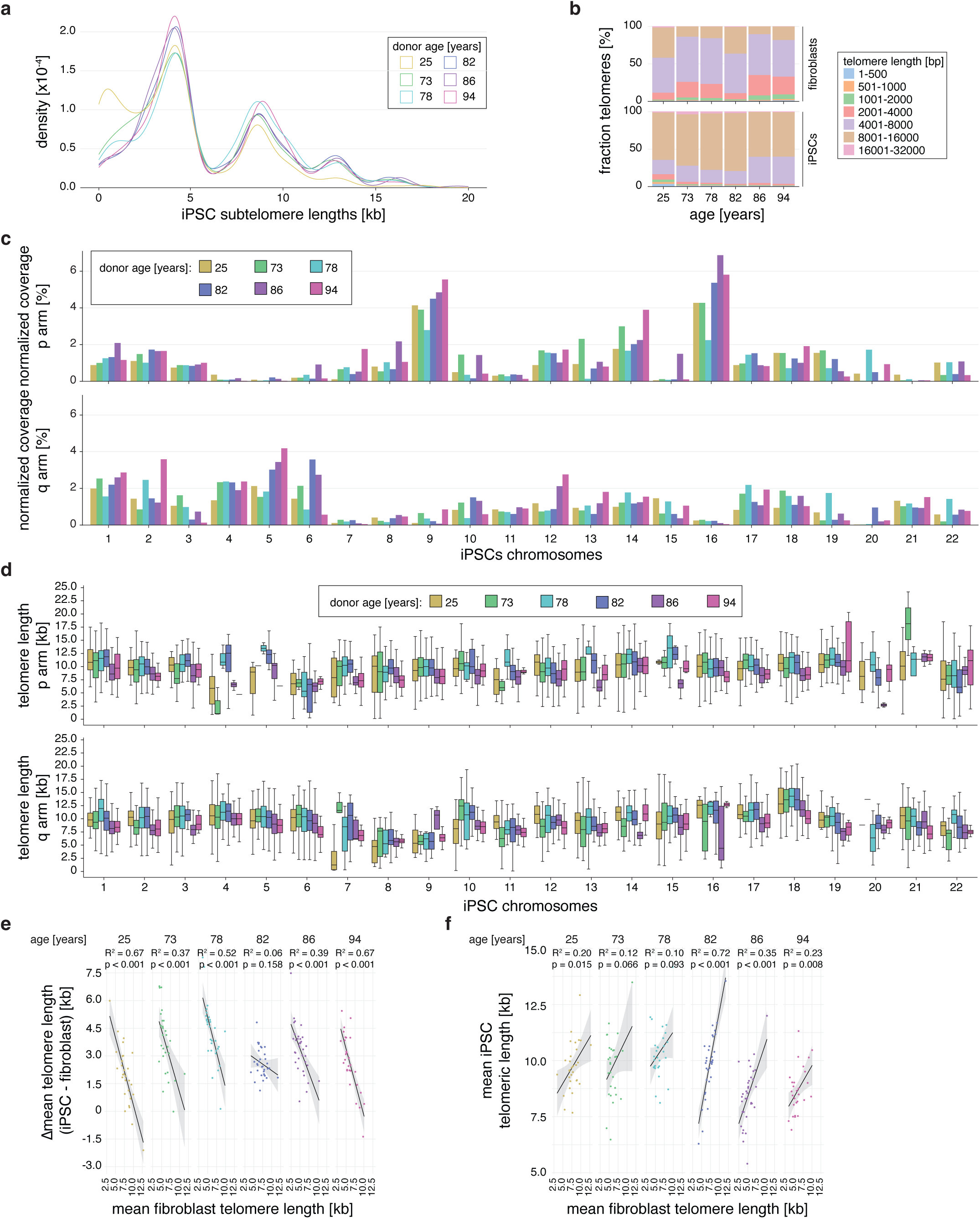
Telo-seq on matched iPSCs. **a,** Subtelomeric length density of matched induced pluripotent stem cells (iPSC). Subtelomeric length is given in kilobase (kb). **b,** Bar graph showing the percentage of binned telomere length in donor-derived matched fibroblasts (top) and iPSC (bottom). **c,** Bar graph of the normalized coverage of telomeric reads per chromosome arm of matched iPSCs of the indicated donor age in percent. **d,** Boxplot of chromosome arm-specific telomere length in kb of matched iPSCs with the indicated donor age. The middle line represents median, box the interquartile range (IQR) and whiskers the 1.5-fold IQR. **e,** Linear regression analysis of the difference between the mean chromosome arm-specific telomere length of matched iPSC and fibroblasts against the mean chromosome arm-specific telomere length in fibroblasts in kb. Black line represents best fit and grey area the 95% confidence interval. Only chromosome arms with at least 20 reads per sample were included. **f,** Linear regression analysis of the mean chromosome arm-specific telomere length of matched iPSC against fibroblasts in kb. Black line represents best fit and grey area the 95% confidence interval. Only chromosome arms with at least 20 reads per sample were included.

**Fig. S6.**
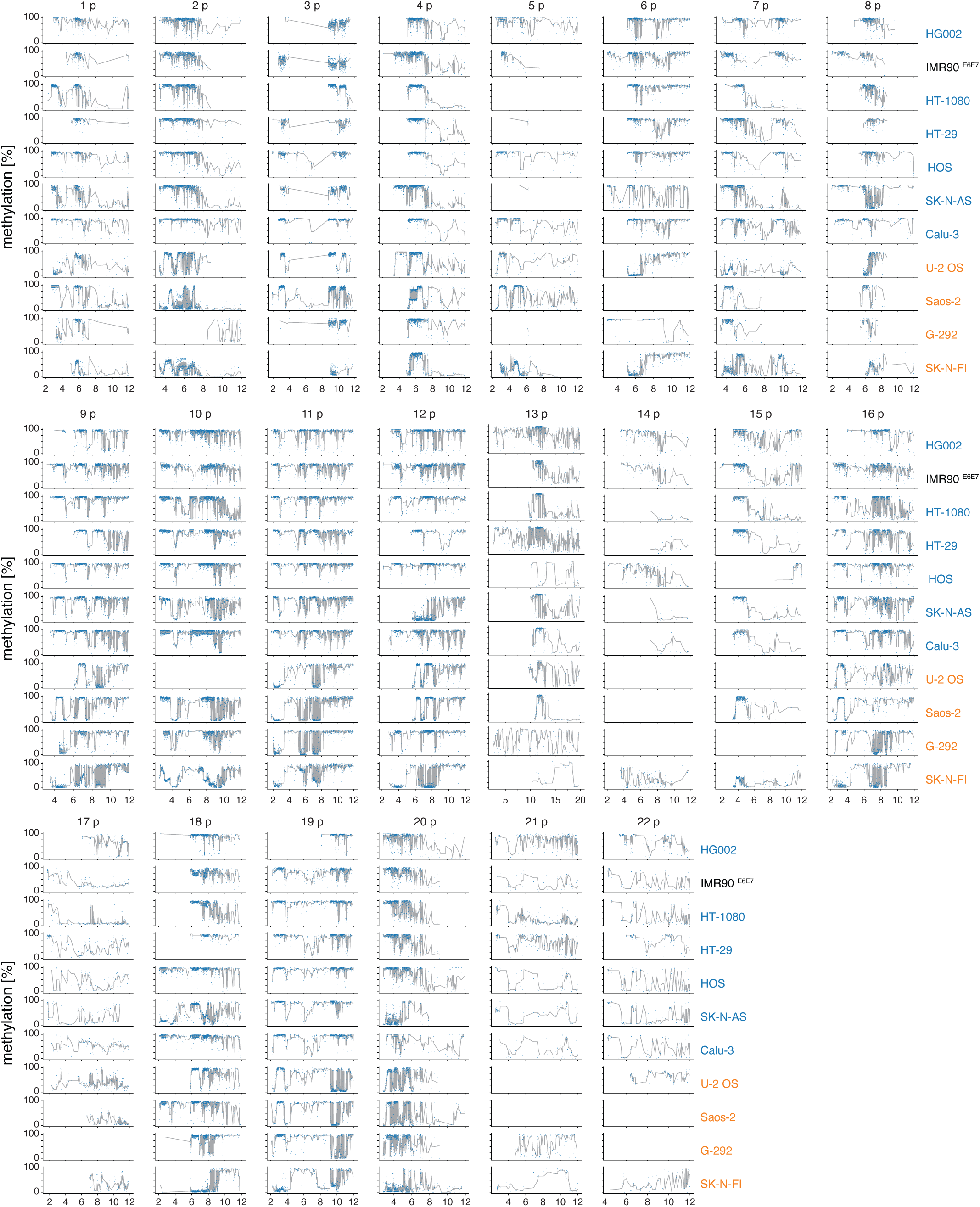
Subtelomeric methylation p chromosome arms. Line graphs of subtelomeric CpG islands closest to telomeres. The percentage of methylation is shown for indicated cell lines and chromosome arms at their kilobase position. Individual CpG sites are represented by blue points and the rolling methylation median is shown as a gray line with a window size of 4. For IMR90 ^E6E7^ the methylation at population doubling 66.2 is shown. Cell lines are color-coded according to telomere maintenance mechanism (TMM); cells with no TMM are in black, telomerase-positive cells in blue and ALT-positive cells in orange. Only data for chromosomes with at least 10 reads per arm are shown.

**Fig. S7.**
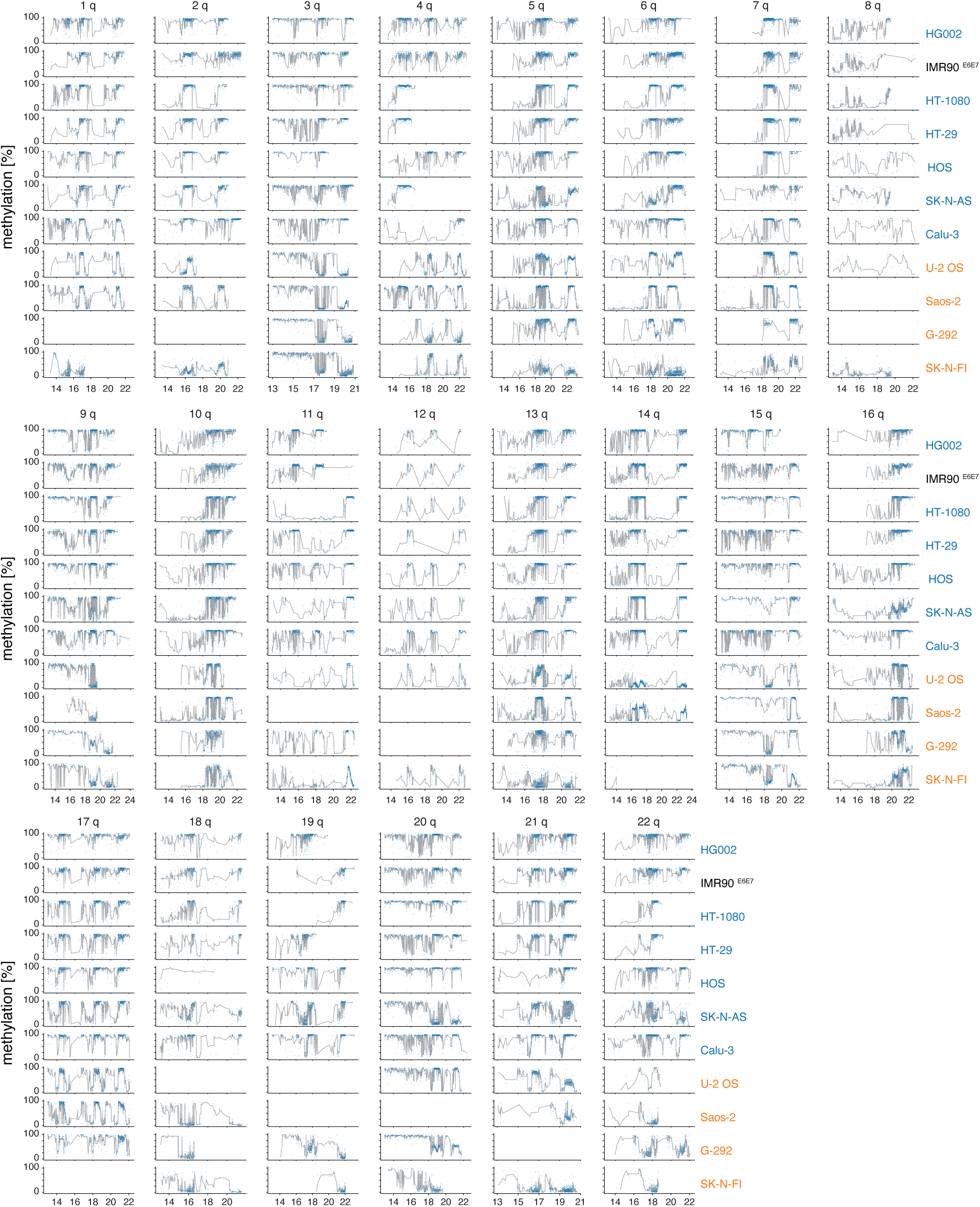
Subtelomeric methylation q chromosome arms. Line graphs of subtelomeric CpG islands closest to telomeres. The percentage of methylation is shown for indicated cell lines and chromosome arms at their kilobase position. Individual CpG sites are represented by blue points and the rolling methylation median is shown as a gray line with a window size of 4. For IMR90 ^E6E7^ the methylation at population doubling 66.2 is shown. Cell lines are color-coded according to telomere maintenance mechanism (TMM); cells with no TMM are in black, telomerase-positive cells in blue and ALT-positive cells in orange. Only data for chromosomes with at least 10 reads per arm are shown.

